# Single-cell phenotypic plasticity modulates social behaviour in *Dictyostelium discoideum*

**DOI:** 10.1101/2022.10.10.511564

**Authors:** Mathieu Forget, Sandrine Adiba, Silvia De Monte

## Abstract

In *Dictyostelium* chimeras, strains social behaviour is defined based on their relative representation in the spores – the reproductive cells resulting from development – referred to as spore bias. Some strains, called ‘cheaters’, display systematically positive spore bias in chimeras and are considered a threat to the evolutionary stability of multicellular organization. The selective advantage gained by cheaters is indeed predicted to undermine collective functions whenever social behaviours are genetically determined. However, genotypes are not the only determinant of spore bias, and the relative role of genetic and plastic phenotypic differences in strains evolutionary success is unclear.

Here, we control phenotypic heterogeneity by harvesting cells in different growth phases, and study the effects of plastic variation on spore bias in chimeras composed of isogenic or genetically different populations. Spore bias is shown to depend both on growth phase and on population composition, and to be negatively correlated to the fraction of ‘loners’, *i*.*e*. cells that do not join aggregates. We examined several single-cell mechanical properties that are expected to affect aggregation efficiency, and found that variations in the fraction of slowly moving cells with growth phase may explain why earlier cultures appear to be underrepresented in the spores. The involvement of a go-or-grow mechanism during cell aggregation is also consistent with known variations of cell-cycle phase distribution during population growth. We confirm the expected ubiquity of growth-phase induced spore bias variation by showing that it is not negligible in genetic chimeras, and can even reverse the classification of a strain’s social behaviour. These results suggest that aggregation can provide an efficient ‘lottery’ system to harness the evolutionary spread of cheaters.

## Introduction

A recognized function of multicellular organization is division of labour, emblematically represented by somatic cells, whose death contributes to the reproductive success of the germline. Such extreme differences in the fate of cells that belong to the same multicellular structure are also found in a number of unicellular organisms – both prokaryotes and eukaryotes – that have independently evolved the capacity of generating multicellular, differentiated structures by aggregation of formerly free-living cells. The most spectacular examples of such aggregative multicellular life cycles are provided by cellular slime moulds, among which *Dictyostelium discoideum* has become a model organism for evolutionary biology (Strassmann and Queller, 2011; Forget et al., 2021).

When they run out of food, cells of *D. discoideum* converge to form multicellular aggregates, that subsequently differentiate in two main terminal fates. One type dies forming the stalk that lifts the other – the spores – above ground and favours their dispersion (Raper, 1940; Smith et al., 2014). In paradigmatic multicellular organisms, where the body derives from the clonal growth of the zygote, the coexistence of germ and somatic cells is facilitated by their genetic uniformity. In aggregative microbes, conversely, where barriers to coaggregation of cells with different genetic background are weaker, multicellular structures often harbour different lineages (Fortunato et al., 2003; Gilbert et al., 2007; Sathe et al., 2013). Conflicts in reproductive investment among genetically diverse cells are expected to threaten collective functions, and to have been particularly acute at the transition to multicellular organization (Rainey and De Monte, 2014). In *D. discoideum*, such conflicts are evidenced by compar-ing, for one of the co-aggregating types, the fraction of cells that are passed on to the following generation (the spores) to the fraction of cells that were present before aggregation (that sets the null expectation for spore pool composition, in the absence of differential reproductive success). ‘Spore bias’ is thus used to identify strains that have qualitatively different social behaviour (Kuzdzal-Fick et al., 2010, 2011; Gilbert et al., 2007; Buttery et al., 2009): cheaters increase their representation in the following generation with respect to cooperators, who on the contrary reduce it.

Theory predicts that, all else being equal, a positive spore bias results in an increase in the frequency of cheaters across multiple social cycles of aggregationdevelopment-dispersal. Hence, in the absence of mechanisms that produce positive assortment between cells with different social investment, cheaters should prevail on the evolutionary time scale, in what is known as the ‘tragedy of the commons’ (Hardin, 1968; Rankin et al., 2007). This conclusion is based on two assumptions: first, that spore bias profiles (spore bias as a function of frequency of the focal strain) are genetically determined, so that they are maintained over the time-scale where cheaters and cooperators strains compete; second, that the spore bias of cheater strains is positive for any frequency of the other type it interacts with.

These assumptions are violated in some cases at least. It is for instance well known that the probability of forming spores, hence spore bias, is affected by phenotypic variation also when cells that co-aggregate are isogenic. For instance, cells have been shown to form spores with different propensity, depending on nutritional history (Leach et al., 1973), cell cycle phase (Zada-Hames and Ashworth, 1978; McDonald and Durston, 1984; Huang et al., 1997; Araki et al., 1994; Azhar et al., 2001; Gruenheit et al., 2018), and duration of starvation (KuzdzalFick et al., 2010) among others (reviewed in (Chattwood and Thompson, 2011) and Forget et al. (2021)). Therefore, spore bias is not exclusively a function of the genetic background of cells, but also of the environment –biotic and abiotic– and of cell physiology. Moreover, several studies of pair-wise chimeras showed that the sign of the spore bias can change with the frequency of the focal type in *D. discoideum* (Madgwick et al., 2018) or in closely related species (Sathe and Nanjundiah, 2018), thus leading to the prediction that cheaters would not exclude cooperators on the evolutionary time scale. These observations challenge the idea that ‘cheater’ genotypes always pose an actual problem to the evolution of multicellular function. However, the extent to which phenotypic variation modifies genetically-established spore bias profiles is unclear. Indeed, experiments have dealt separately with co-aggregation of populations whose phenotypic differences were of plastic or genetic origin (Jang and Gomer, 2011; Strassmann and Queller, 2011; Forget et al., 2021). Despite recent advances in understanding gene regulation in the course of development (Gruenheit et al., 2018; Antolović et al., 2019; Noh et al., 2020; Katoh-Kurasawa et al., 2021), the relation between gene expression and general organizing principles that have been proposed to link cell-level properties to spore bias in chimeras is still largely uncharted.

We have designed an assay to study within a common framework how plastic variation influences social behaviour in binary mixtures of cells with the same or different genetic background. The phenotypic state of cells is (continuously, in principle) modulated by changing the growth phase of cultures when starvation-induced aggregation begins. On the one hand, we can thus affect the nutritional state of cells – a factor that was suggested to be primordial in defining cell fate (Leach et al., 1973; Thompson and Kay, 2000; Zahavi et al., 2018). On the other hand, differences in aggregation timing seem to be more physiological than those imposed by well-distinct culture conditions, even though the amplitude of the difference is enhanced here for effects to be measurable despite unavoidable experimental variability. The ensuing ‘chronochimeras’, obtained by co-aggregation of cells harvested at different growth phases, can be realized both when the two populations have the same or a different genotype, so that cell phenotypic variation is driven by both plastic and genetic differences.

First, we show that, for two populations with the same genotype mixed in various proportions, the time of harvesting affects quantitatively, and sometimes even qualitatively, the spore bias profile, to an extent comparable to that observed when mixing genetically different populations. We next address how physiological variation acquired in the course of growth, which defines the state of the populations at the beginning of aggregation, gives rise to spore bias. Differences in the proportion of non-aggregated cells suggest that early-established biases can impact reproductive success – coherently with previous studies on ‘loner’ cells (Dubravcic et al., 2014; Tarnita et al., 2015; Martínez-García and Tarnita, 2016, 2018; Rossine et al., 2020) – independently of other effects that they may have during multicellular development. The observation that singlecell physical properties change during growth, moreover, supports the idea that cell self-organization in the very first phases of the multicellular cycle may impact evolutionarily relevant biases in more general circumstances, as indicated by numerical models (Garcia et al., 2014; Joshi et al., 2017; Forget et al., 2022). We verify that, according to this hypothesis, spore bias modulation by a change in aggregation timing also occurs when mixing genetically distinct strains. Not only phenotypic effects combine with genetic differences in determining the social behaviour of cells, but – by modifying the frequency-dependence of spore bias – they can change the qualitative nature of the ensuing evolutionary dynamics. Our results confirm that understanding how cells self-organize into aggregates can be as important as deciphering multicellular development for predicting the evolution of social strategies in facultatively multicellular microbes. Moreover, they suggest that simple and general phenotypic differences, such as in cell motility, could translate a multiplicity of molecular mechanisms in their evolutionary effects – something that may illuminate on the emergence of aggregative multicellularity from ancestral unicellular microbes.

## 1. Results

### Chronochimeras

We designed an experimental protocol to examine the combined effects of different sources of cell-cell diversity on the social outcome of co-aggregation of two *Dictyostelium* populations, at least one of which is the axenic lab strain AX3 (Loomis, 1971). Plastic variation in cell phenotype at the moment of aggregation was induced by changing the phase of vegetative growth where cultures were harvested. Populations progression in the growth cycle is indeed known to induce changes in single-cell properties mediated by the accumulation of secreted factors (see Gomer et al. (2011) for a review). For instance, cells harvested at low density are round whereas cells harvested at higher density have multiple pseudopodia, hence possibly different mechanical interactions with the environment and other cells (Yuen et al., 1995). Parameters that correlate with spore bias, e.g. cell-cycle phase distribution and nutritional status (Leach et al., 1973; Azhar et al., 2001; Kuzdzal-Fick et al., 2010), are moreover expected to change as a growing population moves from exponential to stationary phase (Soll et al., 1976; Gomer et al., 2011). A same initial density of cells was grown in standard culture medium for different time intervals. In order to approach natural conditions, minimize the presence of polynucleated cells (Pollitt and Insall, 2008), and avoid drastic changes in their interaction with surfaces, cells were grown in unshaken culture flasks for periods ranging from 24 to 92 hours before harvesting (see Methods and growth curves in Fig. S1). At the beginning of aggregation, cultures were thus in one of four growth phases: Early, Mid, Late Exponential (EE, ME, LE, respectively) and Early Stationary phase (ES). We expect that the phenotypic properties of cells change continuously from one phase to the other, so that the time of harvesting acts as a tunable control parameter.

As illustrated schematically in Fig. 1, two populations harvested at different phases of growth were starved by replacing the growth medium with buffer, then mixed in different proportions at a reference time *t* = 0 h. We call *chrono-chimeras* such binary mixes, whether they belong to the same or different strains. In order to count individual cells belonging to different populations, we transformed AX3 strains with plasmids that bear a fluorescent protein gene (GFP or RFP, see Methods). This fluorescent labelling is maintained in the course of the multicellular cycle, allowing us to quantify, by flow cytometry (see Methods), the proportion of cells of a focal type before aggregation (*f*), and among the spores (*f*_*S*_).

**Figure 1:**
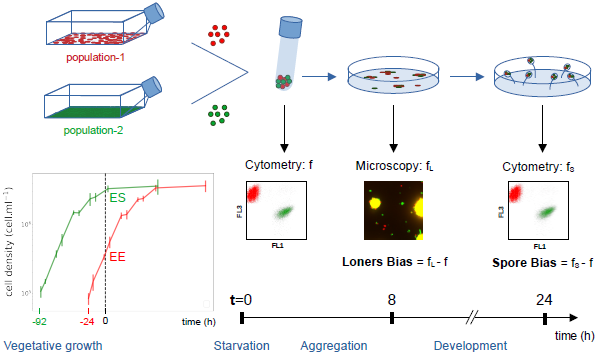
Schematic representation of the experimental protocol to produce chrono-chimeras. Cell populations carrying green or red fluorescent markers were harvested at a specific phase of their logistic growth. Cultures were started at different discrete times before the beginning of the experiment. When the culture was started 24 hours before, cells were called Early Exponential (EE), and they were called Mid-Exponential (ME), Late Exponential (LE), or Early Stationary (ES) when they had been cultured for 46, 68 or 92 hours, respectively. Figure S1 illustrates where in the culture’s growth curve these time are located. As detailed in the Methods, different measures were realized in order to characterize the way biases got established in the course of the multicellular cycle, which was triggered at time *t* = 0 by cell starvation and plating on Petri dishes covered with Phytagel. Before the start of aggregation, the fraction *f* of cells of one population was measured by flow cytometry. Similarly, the fraction *f*_*S*_ of the same population in the spores was performed after completion of development, 24 hours later. Time lapse movies of the aggregation were recorded on an inverted microscope, allowing to count the proportion *f*_*L*_ of each population within the fraction of cells that remained outside aggregates (the so-called ‘loners’). Moreover, measures of single cell properties (discussed later in the text) were realized at *t* = 0 in order to connect initial phenotypic variability to realized biases.

The social cycle is started by plating a binary cell mix on Phytagel. The aggregation (about 8 hours long) is followed by multicellular development, which results in the formation of mature fruiting bodies. Spores were collected by washing whole dishes after 24h. The *spore bias of the focal strain* (specified for every assay) could be thus quantified as the deviation *f*_*S*_ *− f* of its proportion in the spores from that in the initial mix.

### Variable frequency-dependent biases in isogenic chronochimeras

First, we quantified biases in spore production induced by growth phase differences in isogenic chrono-chimeras. Frequency-dependence in spore bias is the basis for inferring evolutionary trajectories, as it connects the composition of the population at the beginning of a social cycle – *i*.*e*. the onset of aggregation – to its composition at the beginning of the following. Hence, we assessed several initial compositions of the binary mixes, so as to derive spore bias as a function of the fraction of cells of the focal type *f* (referred to, in the following, as *spore bias profile*).

Figure 2 displays the spore bias profiles when populations harvested at different stages of vegetative growth (ME,LE,SE), labeled with GFP, are mixed with a RFP-labeled reference EE population. Insertion of different fluorescent proteins introduced a reproducible intrinsic bias, which was used to correct the measures by a frequency-dependent term identical for all chrono-chimeras (see Methods).

**Figure 2:**
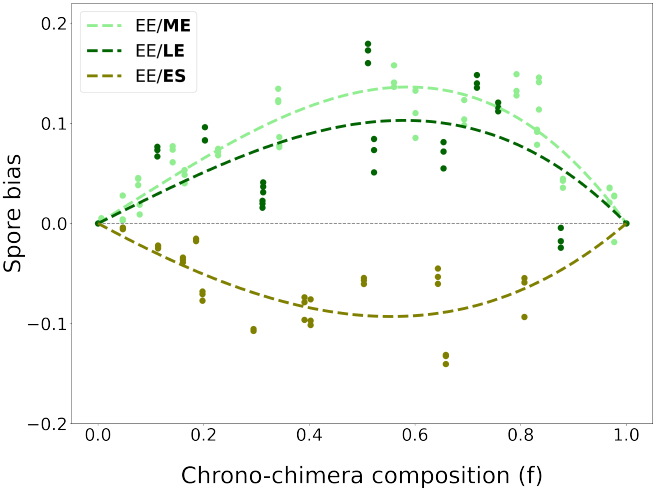
Spore bias profiles depend on growth phase-induced phenotypic variation. Corrected (see Methods) spore bias for a focal AX3-GFP population harvested in different growth phases (indicated in bold in the legend), mixed to a reference AX3-RFP EE population, as as function of the proportion *f* of the focal strain. Measures of spore bias were realized in three replicates (same initial mix plated on three different plates) and the error bars indicate the standard deviation of the spore measure realized 24*h* later. The observations have been interpolated (dashed lines) with a third degree polynomial constrained with *f* = 0 and *f* = 1.

Our observations indicate that phenotypic differences induced by the phase of population growth at the beginning of aggregation bias spore production. The spore bias profiles are frequency-dependent and resemble in shape and magnitude those observed in genetic chimeras. Even though we never observed spore bias profiles that changes sign at intermediate frequencies (Sathe and Nanjundiah, 2018; Madgwick et al., 2018), it is possible that this may manifest if we sampled more extensively the growth curve.

ME and LE populations are reproducibly associated with a positive spore bias when co-aggregating with EE populations (as was previously observed when mixing cultures that had been starved for different periods (Kuzdzal-Fick et al., 2010)). However, such advantage appears to wane for older ES cultures, that can display a negative bias. Different isogenic populations would be therefore alternatively classified as cheaters or cooperators depending on their ecological history at the onset of aggregation.

### Efficiency of aggregation links phenotypic variation to spore bias

Growth phase at the onset of aggregation is thus able, together with other documented sources of plastic phenotypic variation (Leach et al., 1973; ZadaHames and Ashworth, 1978; Azhar et al., 2001; Hiraoka et al., 2020), to affect the probability that a cell will turn into a spore. How early phenotypic heterogeneity results in biases that manifest themselves many hours later is generally unknown. Indeed, if genetic differences are maintained on the time scale of the social cycle, and beyond, differences associated to growth phase may be readily erased after aggregation, when cell signalling – common to all cells in the chimera – drives cell differentiation.

If growth-phase induced spore biases were chiefly due to processes acting within multicellular aggregates, one should suppose that social interactions were primed by phenotypic differences present several hours before the aggregation is completed. Such long-lasting phenotypic imprinting may then result in sorting within the slug, as commonly observed when mixing different genotypes (Ennis et al., 2000; McDonald and Durston, 1984; Houle et al., 1989; Escalante et al., 1997). In our case, however, no noticeable segregation or sorting was observed either in the mound stage or during slug migration (Fig. S4 A and B), suggesting that social, strategic interactions within the aggregates may not be the main factor determining spore bias differences in chronochimeras.

Alternatively (or additionally) biases may get established early enough, so that initial differences among co-aggregating cells matter, even if these differences are inconsequential at later developmental stages. This hypothesis was proposed in models that stressed the evolutionary relevance of cells that remain outside aggregates, called ‘loners’ (Dubravcic et al., 2014; Tarnita et al., 2015; Martínez-García and Tarnita, 2016, 2018; Rossine et al., 2020). Different strains were shown to leave a different proportion of non-aggregated cells (Dubravcic et al., 2014; Rossine et al., 2020), however spore bias was not directly assessed in those experiments.

We therefore considered whether the spore biases observed in isogenic chronochimeras could reflect a disproportional representation of phenotypically different populations within aggregates, thus also in the fraction of non-aggregated cells. We measured, at a time when mounds were completely formed (about 8 hours into the social cycle, see Fig. 1), the loner bias. This was quantified (see Methods) in isogenic chrono-chimeras by subtracting the proportion of the focal population in the pool of loners to its expected proportion, that is the initial fraction *f ∼* 50%. We decided to focus on this relative measure because it is very complicated to count the absolute number of loners over a whole field of aggregation, and moreover the loner bias compares directly to the spore bias. Figure 3 shows that the loner bias is negatively correlated to spore bias when chrono-chimeras with different growth differences are taken into account (Pearson correlation coefficient= *−*0.58, *p* = 0.01). It should be noted, however, that the loner bias of ES populations varied when the experiment was repeated, possibly reflecting the difficulty of precisely controlling the conditions that cells meet at the entry in stationary phase.

**Figure 3:**
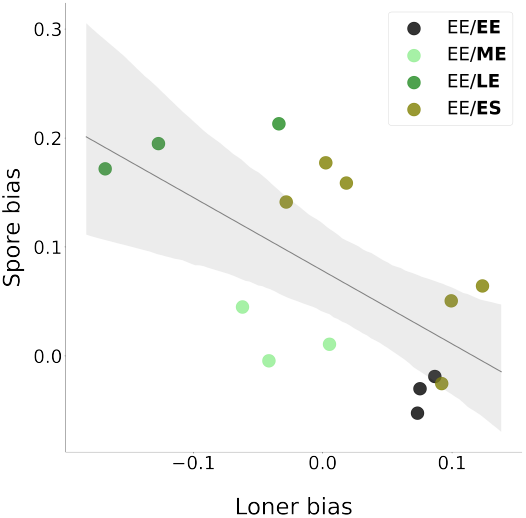
Spore bias negatively correlates to loner bias. Corrected spore bias (see main text and Methods) as a function of loner bias in isogenic chrono-chimeras composed of the strain AX3-GFP with *f ∼* 50% of a reference AX3-RFP EE population. The growth phase of the focal population (AX3-GFP) is indicated in bold in the legend.

Cultures that leave more cells as loners, are thus under-represented in the pool of spores, as would be expected in the absence of strong developmental biases. A substantial part of spore bias variation might then be attributed to cells being more or less efficient in aggregating, depending on their growth phase. In particular, it appears that aggregation is maximised at intermediate times during logistic growth, while cultures that are close to the stationary phase tend to leave more cells behind.

### A go-or-grow single-cell mechanism may link growth phase to aggregation efficiency

In order to understand how initial phenotypic differences lead to the observed biases in aggregation efficiency, we looked for relevant single-cell parameters that might operate early in the social cycle. Differences in sensitivity to an external, diffusing signal was proposed to underpin differential aggregation propensity, and a mathematical model confirmed that this mechanism can result in variable proportions of loner cells (Rossine et al., 2020). If it is biologically reasonable to assume that signalling gets affected by growth phase, and that differences in the perception of signals may last long enough to cause differential aggregation, the specific molecules involved in this process have not yet been identified.

Another possible – and by no means alternative – explanation of loner biases is that different aggregation propensity stems from changes, along the culture growth, of cell mechanical properties. Numerical models for cell selforganization into groups indeed show that differences in cell-to-cell adhesion or velocity can lead to loner biases and be potentially involved in the evolution of aggregative multicellularity (Garcia et al., 2014, 2015; Miele and De Monte, 2021; Forget et al., 2022). We thus characterized the variation during vegetative growth of three parameters that are involved in single-cell behaviour and in short-range cell-cell interactions.

First, we considered surface and cell-cell adhesion (that can also encompass systems evolved for self-recognition (Hirose et al., 2011)). The former was quantified (see Methods) by measuring the fraction of cells that adhere to a culture vial at low cell density, so that direct interactions among cells should be negligible and our measures pertain to traits of individual cells. Cell-cell adhesion was quantified by measuring the percentage of cells that formed multicellular clusters in shaken cultures (see Methods). Cell adhesion was found to change gradually during demographic growth. Cell-substrate adhesion significantly increases in the course of vegetative growth (Fig. 4 A). Cell-cell adhesion, on the contrary, significantly decreases as a population ages (Fig. 4 B). Studies on the role of differential adhesion in the evolution of social behaviour focused on cell-cell interactions, and predict that less adhesive cells are found more often among the loners (Garcia et al., 2015), which is not what we observe. Another expectation that is not met in chrono-chimeras (Fig. S4 A) is that if cells were able to recognise the internal state, differential cell adhesion would, like for kin recognition, induce segregation of the two co-aggregating populations into aggregates that are mainly composed by one or the other type. We therefore presume that, beyond ‘social’, cell-cell contacts, cell-substratum adhesion plays a key role in establishing social behaviour, as also supported by recent directed evolution experiments (Adiba et al., 2022).

**Figure 4:**
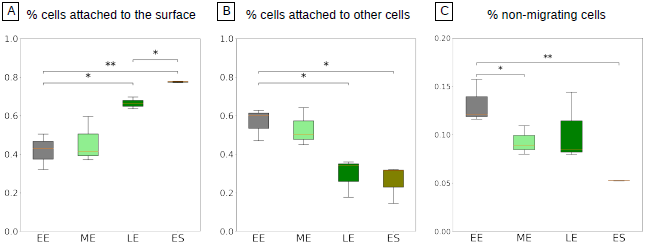
Variation of cell adhesion and motility properties in the course of vegetative growth of AX3 populations. Cell-substrate adhesion increases (A), cell-cell adhesion decreases (B) and the proportion of non-migrating cells (see Fig. 5) decreases as a population ages (C). In each assay, three replicates were performed for each condition. **\***: Student test p-value *<* 0.05 **\*\***: Student test p-value *<* 0.005.

Then, we addressed single-cell motility. As amoebae crawl by extending pseudopods (Kessin, 2001), variations in adhesion to the substratum may alter the probability of aggregation by altering the speed of displacement. This hypothesis is supported by numerical models for binary mixes of self-propelled particles, that showed that differential motility can result into assortment within the aggregated phase (Forget et al., 2022; Kolb and Klotsa, 2020; S. Punla et al., 2022).Moreover, cell motility has been recently invoked as the basis of differentiation biases observed in cells with different intracellular ATP concentration (Hiraoka et al., 2020, 2022). We measured individual cell motility in populations harvested in EE, ME, LE, and ES phase.

Individual cell motility can be studied by tracking single cells in diluted cultures, so that encounters are rare. We analyzed a large number of cell trajectories (*∼* 600 for each condition) in populations harvested at different growth phases and diluted before realizing time-lapse movies (see Methods). Cell position was measured every 30 seconds in the course of 1 hour. The slope of individual Mean Square Displacement (MSD) as a function of time lag Δ*t* (Fig. 5 A and Fig. S5) reveals the co-existence of two classes of cells with markedly different motility. Figure 5 D shows that part of the cells barely move during the experiment (‘non-migrating cells’, red), while the others efficiently crawl on the substratum (‘migrating-cells’, blue). Differences between these classes, quantified for instance by their total displacement, are highly significant (Mann–Whitney U test p-values *<* 0.0005, Fig. 5 C and Fig. S6). A bimodal distribution of cell motility is consistent with a previous analysis of a small number (*∼* 40 cells) of *Dictyostelium* AX2-cells just before starvation (Goury-Sistla et al., 2012)). Cell speed within these two motility classes is not significantly different in different growth phases (total displacement of migrating cells was compared with a mixed effects model ANOVA, p-value=0.4691, Fig. S6). However, growth phase alters population partitioning between slow and fast cells, the percentage of low-motility cells decreasing as cell culture ages (Figure 4 C). By comparing these results to variations in adhesion, we can speculate that cells belonging to the slower class also adhere less to the surface and are more likely to end up as loners. A progressive decrease in the fraction of non-migrating cells, which are more likely to remain outside aggregates, could thus explain why ME and LE cultures tend to have a positive bias relative to EE/EE chrono-chimeras. However, in order to explain the decrease of the bias observed in EE/ES chrono-chimeras, other yet uncharacterized mechanisms should be identified.

**Figure 5:**
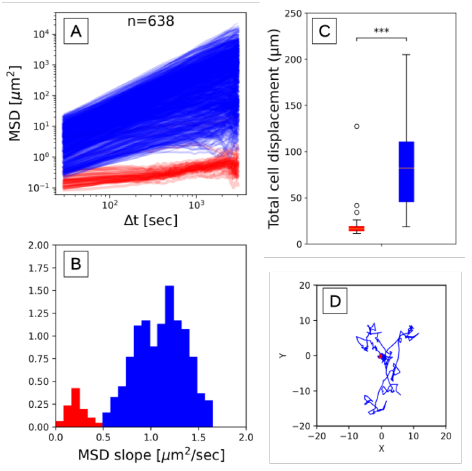
AX3 populations display a bimodal distribution of cell motility. **A:** Individual mean square displacement as a function of time lag (Δ*t*). **B:** Distribution of the initial rate of increase of the MSD (slope of the log MSD vs log Δ*t*, Δ*t <* 150 secs). Cells were clustered into two motility classes (indicated in all panels in red and blue respectively), where such slope was below or above the threshold value 0.5. **C:** Total displacement of cells from these two classes in the course of 1 hour, showing that the initial slope is a good proxy for how much cells displace (**\*\*\***: Mann–Whitney U test p-value *<* 0.0005.) **D:** Typical cell trajectories from the two motility classes with their origins brought to a common point, illustrating the difference in motility. Results are shown for one LE population (see Figs. S5 and S6 for all replicates and growth phases).

The observed variations in cell motility and surface adhesion are consistent, as we discuss later, with a change during population growth of cells distribution among different phases of the cell cycle. Older cultures are indeed enriched in cells that have stopped their progression through the cell cycle and are blocked in G2 (Soll et al., 1976). It is known that G2 cells have a higher chance of becoming spores (Zada-Hames and Ashworth, 1978; Gomer and Ammann, 1996; McDonald and Durston, 1984; Gruenheit et al., 2018). Our results suggest that, by the relation of cell cycle phase and motility, aggregation efficiency mediates between the initial cell cycle phase differences and spore bias. Like the go-orgrow hypothesis (Giese et al., 1996), cells that are dividing may make up the low motility subpopulation. Rearrangement of the cytoskeleton during mitosis indeed would cause cells to detach from the substratum (Nagasaki et al., 2002; Plak et al., 2014) and therefore reduce their migration efficiency, underpinning the concomitant increase in surface adherence and motility during culture growth.

The go-or-grow mechanism is independent of possible later effects of cell cycle phase on differentiation within the multicellular aggregates. As suggested by theoretical models (Forget et al., 2022; Kolb and Klotsa, 2020; S. Punla et al., 2022), it may thus be relevant more broadly, whenever two cell types with different mechanical properties co-aggregate. In particular, we expect that growth-phase induced biases manifest also when the populations that are mixed belong to distinct strains. They could, however, be negligible with respect to genetically-induced biases, thus making the spore bias profile largely independent of the specific experimental settings, and thus predictive of long-term competition among strains.

### Growth phase affects social behaviour also in genetic chronochimeras

We considered chrono-chimeras obtained by mixing strains whose synchronous co-aggregation resulted in differential spore production. We compared spore bias profiles in three conditions: when the two strains were both harvested in the ME phase (see Fig. S1 for the growth curves of different strains), when the focal strain was in EE and the other in LE, and vice-versa. The first condition corresponds to standard settings, where the two populations are harvested at the same time, and is used to set the ‘baseline’ spore bias profile. If growth phase had the same effect as for isogenic chronochimeras, the latter two conditions would induce changes of the profile in opposite directions. Quantification of these effects allows to determine if and when such changes are comparable to those induced by distinct genetic backgrounds.

In line with the hypothesis that adhesion plays a key role in determining the efficiency of aggregation, we started examining chimeras composed of strains with highly divergent cell-substratum adhesion. The AX3-Bottom strain was evolved from the ancestral AX3 strain by imposing selection for increased adherence to a culture vial, and has a strong spore bias relative to the ancestor (Adiba et al., 2022). Differences between AX3 and AX3-Bottom are not known at the genotypic level, but these strains are expected to diverge chiefly for the phenotype that was under selection (single cell adhesion to the substratum). Compared to the ‘baseline’ profile, spore bias of the focal strain increased or decreased for all frequencies, depending on whether it is harvested later or earlier (Fig. S6 A). Similar results are obtained when the more advanced population is harvested in ME phase rather than in LE phase (Fig. S6 B). Spore bias is thus concomitantly affected by plastic variation and phenotypic differences resulting from selection on single-cell adhesion. Even though growth differences modify the bias, however, they are not sufficient to alter the qualitative classification of social behaviour. The ancestor strain keeps being classified as a cheater, so that it is predicted to outcompete the evolved strain over multiple rounds of aggregation and dispersal.

A similar growth phase-dependent variation in the bias was obtained in chrono-chimeras where the reference strain AX3 was mixed with another strain derived from an AX3 ancestor, the well-known cheater strain chtA (Ennis et al., 2000). ChtA displays the most extreme and disruptive form of selfish behaviour, obligate cheating: it is unable to form spores when developing clonally, but induces its ancestor to differentiate into stalk. Consistent with this classification, we found that the AX3 strain had a negative bias when harvested at the same time as chtA (Fig. 6 A). Such bias tends to increase when AX3 is harvested in EE phase and chtA in LE phase, but is reduced to almost zero for the reversed growth phase relation (when AX3 is in high proportion in the chrono-chimera), confirming again that populations in different growth states can produce variable contributions to the spore pool.

**Figure 6:**
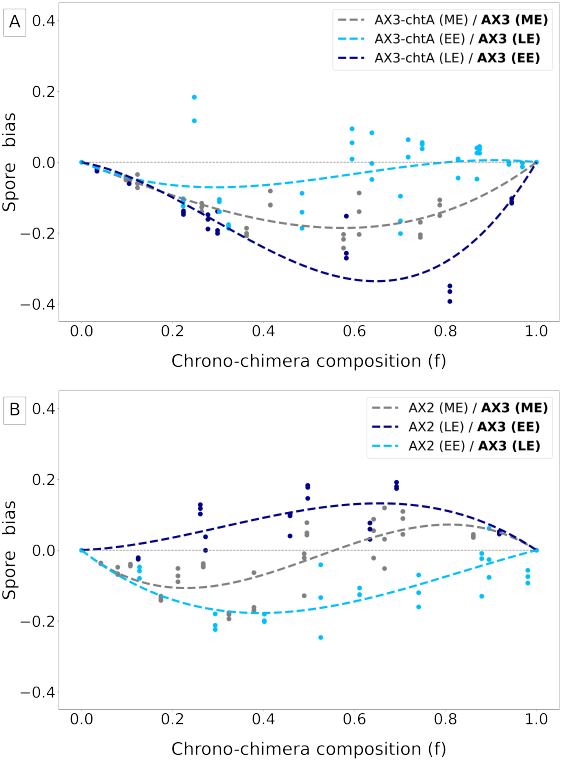
Social behaviour in chimeras depends on populations growth phase at the onset of aggregation. Spore bias measured in two different genetic chimeras, using a population of AX3-RFP cells as the focal population. **A:** chrono-chimeras composed of AX3-RFP and chtA cells. **B:** chrono-chimeras composed of AX3-RFP and AX2 cells. Three different cases are considered: strains are grown in co-culture and harvested in ME phase, the focal population is harvested in EE phase and the other in LE phase (dark blue line), or vice-versa (light blue line).

AX3-Bottom and AX3-chtA were obtained by artificial selection (directional selection of single-cell adhesion and mutagenesis followed by screening for strong spore biases, respectively). Correspondingly, they manifest extreme social behaviours. However, in natural conditions co-aggregating strains may not be as phenotypically divergent, having evolved under selective pressures that likely were both weaker and acting on many traits simultaneously. We examined chrono-chimeras for another pairwise combination of strains, where AX3 was mixed with the axenic strain AX2. Both these strains derive from the same natural isolate (NC-4), but have genome-wide differences: AX3 carries a large duplication, corresponding to 608 genes (Sucgang et al., 2011). Despite such large genomic differences, AX2 is not known to display a marked social behaviour with respect to AX3. Indeed, we found that, in the absence of growth phase heterogeneity, AX3 cells co-aggregating with AX2 cells are under-represented in the spores when in low proportion in the chimera, whereas they are associated to a positive spore bias when prevalent (Fig. 6 B).Spore bias profiles that change sign with the frequency of one strain in the mix are commonly observed in natural isolates (Madgwick et al., 2018; Sathe and Nanjundiah, 2018), and are likely to be more representative of interactions in the wild than the previously considered chimeras. In the AX2/AX3 chrono-chimera the focal strain AX3 shifts from behaving like a cheater (when it is harvested in EE while AX2 is in LE) to behaving as a cooperator (in the opposite case).

Taken together, results on chrono-chimeras involving different strains indicate that the effects on social behaviour of growth phase-induced phenotypic differences at the time of aggregation combine with those due to genetic diversity. In some cases, growth phase-induced phenotypic differences can reverse the classification of a strain from cheater to cooperator. The contribution of these two sources of variation to spore bias is not additive. Instead, the direction of change in spore profile as a function of the growth phase of the two cultures depends on the chimeras genetic background, as summarized in Fig. S7.

## 2. Discussion

Division of labour within multicellular structures, whereby different cells take up different tasks, is essential for sustaining collective functions, but is often associated to differences in reproductive success among distinct cell types, e.g. between germ and somatic lineages. Such differences are particularly disruptive when cell heterogeneity is transmitted across generations of the collective association. In aggregative microbes like *Dictyostelium discoideum*, where multicellular groups are formed by gathering previously isolated cells, cell types that are overrepresented in the spore pool have the potential to get, over successive aggregation cycles, progressively enriched in the population. Crucial for this to occur is however a heritable relation between cell genotype and its reproductive success. This is realized when the outcome of social interactions is by and large genetically determined, as assumed by the theory of sociobiology (Strassmann and Queller, 2011).

Several recent studies have started to question the relevance of this assumption for *Dictyostelium* and revealed complex relationships between properties of single cells and their reproductive success. Madgwick et al. (Madgwick et al., 2018) proposed that frequency-dependent spore bias profiles in pairwise chimeras of natural strains are explained by cells adjusting their probability of sporulating as a response to the multicellular context. In this perspective, cheat-ing would result from a ‘strategic’ choice of each cell, influenced for instance by the diffusion of morphogens during multicellular development (Parkinson et al., 2011). Therefore, a same cellular genotype could give rise to multiple possible biases depending on how many and what kind of cells happen to surround a focal cell, giving rise to frequency-dependent spore bias profiles.

Our results show that spore bias profiles depend on the nature of the social partner also in isogenic chrono-chimeras, where recognition of genetic identity is not an issue. Not only the intensity of the bias depends on the composition of the mix (Fig. 2), but a population harvested in early exponential phase can be associated to a positive or a negative spore bias, depending on the growth phase of the co-aggregating population. The correlation between spore and loner bias suggests that biases do not necessarily require ‘negotiations’ involving a multitude of cells, and can get established ahead of multicellular development, as a result of cell self-organization during aggregation.

Similar mechanisms acting at early stages of the social cycle were suggested to underpin the role of loner cells in the evolution of cooperative behaviour (Dubravcic et al., 2014; Tarnita et al., 2015; Rossine et al., 2020). While early models proposed that the probability of aggregation (hence the ensuing spore bias) was determined by strain genotype, additional measures revealed that the situation is more complex, and context-dependence widespread (Dubravcic et al., 2014; Rossine et al., 2020). In the absence of direct measures of spore bias, a mathematical model was used to show that differences in the sensitivity to an aggregation signal induce frequency-dependent loner bias profiles, which can be leveraged for maintaining - over multiple social cycles - coexistence of different, conflicting genotypes (Rossine et al., 2020).

If biases are broadly set before multicellular development, then pre-existing differences in single-cell phenotypic properties can matter as long as they persist throughout the aggregation phase, whether their origin is genetic or plastic. Models for cell aggregation indeed point to the possible role of cell-cell adhesion and motility (Garcia et al., 2014, 2015; Martínez-García and Tarnita, 2016, 2018; Arias Del Angel et al., 2020; Forget et al., 2022) – on top of the chemotactic response to diffusing signals (Rossine et al., 2020) – in establishing loner biases. With this hypothesis in mind, we looked for single-cell mechanical properties that varied with the growth phase of the cell culture.

Cell-surface attachment, cell-cell adhesion and single-cell motility all show related changes during the initial phases of culture growth. We focused in particular on characterizing variation in the distribution of single-cell motility because, on the one hand, motility appears to be regulated by ATP independently from cAMP oscillations (Hiraoka et al., 2022). On the other hand, it can be connected more directly to the observed increase in proportion of aggregated cells as the growth phase advances. When a population ages, indeed, the fraction of actively dividing cells (in the M phase of the cell cycle) is known to decrease (Soll et al., 1976; Zada-Hames and Ashworth, 1978). *Dictyostelium* cells entering cytokinesis tend to round up and to be less adhesive to the substratum (Nagasaki et al., 2002; Plak et al., 2014). As a consequence, they may contribute disproportionately to the pool of non-migrating cells that end up not joining any aggregate, with a mechanism analogous to the “go-or-grow” hypothesis proposed for cancer cells (Giese et al., 1996) and recently applied to *Dictyostelium* motility under hypoxia (Cochet-Escartin et al., 2021). Such a mechanism, moreover, roots the previously reported negative correlation between the fraction of M/S cells and spore production in mechanical processes occurring at the onset of the multicellular cycle, when cell behaviour is least influenced by social interactions (Zada-Hames and Ashworth, 1978; McDonald and Durston, 1984; Huang et al., 1997; Araki et al., 1994; Azhar et al., 2001; Gruenheit et al., 2018).

Mathematical and numerical models showed that heterogeneity in single-cell motility can result in differential partaking of the multicellular organization, with consequences that extend to the evolutionary time scale (Rossine et al., 2020; Miele and De Monte, 2021; Forget et al., 2022). It is therefore possible that some of the principles illustrated by our observations in controlled lab settings may apply more broadly.

Growth phase differences at the moment of starvation are expected to oc-cur in natural populations, where the history of cells during vegetative growth may vary greatly even within a single clone, due to different timing of spore germination, which sets the onset of demographic growth. As cyclic adenosine monophosphate (cAMP), the main signal driving aggregation of *D. discoideum*, diffuses very fast, it seems moreover likely that, in the soil, the aggregation domains of a few centimeters encompass micro-scale variation in biotic and abiotic factors. Here, we have used growth phase as a control parameter to continuously tune the phenotype of cells in a population, and explored large time lags in order to quantify differences with more ease. Whether the hypothesis of synchronous or asynchronous aggregation is closer to natural aggregative cycles would require additional studies in the wild.

Temporal differences are just one possible non-genetic source of cell phenotype variation that affects representation in the pool of spores. Plastic heterogeneity could be caused by both environmental and physiological variation (Leach et al., 1973; Kuzdzal-Fick et al., 2010; Kubohara et al., 2007; Hiraoka et al., 2020). How it gets transmitted across the aggregation phase and through multicellular development is however still unclear. Indeed, given the fast changes in gene expression during *Dictyostelium*’s social cycle (Coates and Harwood, 2001; González-Velasco et al., 2019), phenotypic variation forged during vegetative growth should in principle fade shortly after the beginning of the social cycle. On the contrary, non-aggregated cells are irreversibly excluded from multicellular development, so that differences in aggregation efficiency might explain biases in isogenic populations derived from different sources of plastic variation. The possibility that initial mechanical heterogeneity may compete in more general settings with genetically-established social behaviour will require further exploration.

Single-cell phenotypes can influence aggregation probability irrespective of their genetic or non-genetic origin. However, their underpinning and the way they get transmitted along the multicellular cycle are important to understand the evolution of social behaviours. In particular, the prediction that ‘cheaters’ have a long-term advantage could be upended if short-term measures of repro-ductive success do not carry over to successive generations, so that relevant variation is effectively neutral (Arias Del Angel et al., 2020; Nanjundiah, 2019; Forget et al., 2021). Our results point to a role of unpredictable variation that may be much larger than previously considered when associating a genotype to a social behaviour. Single-cell properties at the moment of aggregation and the derived biases are not only shaped by the genetic identity of a strain, but also by factors that are not under direct genetic control. Such factors have more to do with the ecological history of individual cells and its consequences on cell mechanics than with the genetically-determined behaviour in a multicellular, social context. We can thus speculate that spore bias variability may have similar underpinnings, whatever the origin of phenotypic variation. If such explanation holds true also in natural populations, it may contribute to understand how aggregative multicellular life-cycles persist on evolutionary times despite unavoidable genetic conflicts.

## Acknowledgments

This work has received support under the program “Investissements d’Avenir” launched by the French Government and implemented by ANR with the references ANR–10–LABX–54 MEMOLIFE and ANR–10–IDEX–0001–02 PSL* Uni-versité Paris, Q-life ANR-17-CONV-6150005, and the project ANR-19-CE45-0002 ‘ADHeC’ PSL research University.

## STAR⋆Methods

### Resource availability

#### Lead contact

Further information and requests for resources and reagents should be directed to and will be fulfilled by the Lead Contact, Dr. Mathieu Forget.

#### Materials Availability

Materials generated in this study are available from the Lead Contact with a completed Materials Transfer Agreement.

#### Data and Code Availability

The data and Python scripts generating the figures are available from the Lead Contact on request.

### Key resources table

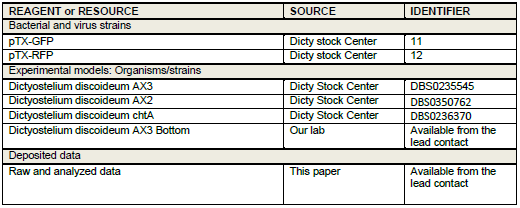

| REAGENT or RESOURCE | SOURCE | IDENTIFIER |
| --- | --- | --- |
| Bacterial and virus strains |  |  |
| pTX-GFP | Dicty stock Center | 11 |
| pTX-RFP | Dicty stock Center | 12 |
| Experimental models: Organisms/strains |  |  |
| Dictyostelium discoideum AX3 | Dicty Stock Center | DBS0235545 |
| Dictyostelium discoideum AX2 | Dicty Stock Center | DBS0350762 |
| Dictyostelium discoideum chtA | Dicty Stock Center | DBS0236370 |
| Dictyostelium discoideum AX3 Bottom | Our lab | Available from the lead contact |
| Deposited data |  |  |
| Raw and analyzed data | This paper | Available from the lead contact |

#### Experimental model and subject details

##### Strains and media

*Dictyostelium discoideum* strains used in this study are AX3 (Dictybase ID: DBS0235545), AX3-chtA (Dictybase ID: DBS0236369), AX2 (donation by Clément Nizak), and the AX3-Bottom line (Adiba et al., 2022). AX3-GFP and AX3-RFP cell lines were obtained by transforming AX3 cells with plasmids pTX-RFP (Dictybase ID: 112) or pTX-GFP (Dictybase ID: 11) (Dubravcic et al., 2014). Both plasmids carry a gene for antibiotic resistance (Gentamicin 418, Sigma-Aldrich: G418). Vegetative growth was started from frozen aliquots thawed every week to prevent the accumulation of undesired mutations due to prolonged culturing. Cells were grown in autoclaved HL5 medium (per L, 35.5 g HL5 from *formedium*, pH=6.7) at 22°C with a concentration of 300 μg mL^*−*1^ Streptomycin. Additional 20 μg mL^*−*1^ G418 were supplemented when growing transformed strains. Pre-cultures were prepared by thawing frozen aliquots and growing in 25 *cm*^2^ TC treated flasks (*CytoOne* CC7682-4825) with 10 mL culture medium for 30 hours, which allows the population to restart and enter the exponential growth phase. Cells were grown in static cultures to limit the risk of impaired cytokinesis as observed in shaken suspension. SorC buffer was prepared with 0.0555 g *CaCl*_2_; 0.55 g *Na*_2_*HPO*_4_ 7*H*_2_*O*; 2 g *KH*_2_*PO*_4_ per Liter.

### Method details

#### Strains growth kinetics

Growth kinetics of AX3-GFP, AX3-RFP, AX3-Bottom, AX3-chtA and AX2 were characterized to estimate the timing of their growth cycle under the experimental conditions (Fig. S1). Pre-cultures were diluted to 10^5^ cells/ml and re-suspended in fresh medium. At each time point, cell density was scored using an hemocytometer. Three replicate cultures of each strain were examined in parallel.

#### Starvation protocol

In order to trigger *Dictyostelium* social cycle, cell populations were starved by washing out the nutrient medium via three successive centrifugations with buffer at 4°C (2000 rpm for 7 min). Cells were kept on ice between successive rounds of centrifugation. After the last centrifugation, the pellet was resuspended in buffer and the cell density adjusted to 2.10^7^ cells mL^*−*1^.

#### Chrono-chimeras preparation

Chrono-chimeras are composed of a mix of starved cells from two populations harvested at different times during vegetative growth, *i*.*e* in different growth phases at the time *t* = 0 when the experiment was begun (Fig. 1). Each population was started from a pre-culture diluted into fresh medium to a density of 10^5^ cells mL^*−*1^. In order for them to attain different growth phases (established based on the growth kinetics displayed in Fig. S1) at *t* = 0, cultures were started a fixed number of hours before the beginning of the experiment: 24h hours for early-exponential phase (EE); 46 hours for mid-exponential phase (ME); 68 hours for late-exponential phase (LE) and 92 hours for early-stationary phase (ES). Beforehand, we made sure that transformed cells used in isogenic chronochimeras had indistinguishable growth curves (Fig. S1) to confirm that populations harvested after the same growth duration were in the same growth phase.

At *t* = 0h (Fig. 1), the two cultures were starved as described in the previous section. Starved cells from the two populations were then mixed so as to attain a target proportion. Deviations from the target proportion sometimes occurred due to fluctuations in dilution. The actual mix composition *f* was thus quantified by measuring the proportion of labelled cells by flow cytometry (Cube8 cytometer, using Forward Scatter (FSC), Side Scatter (SSC), fluorescence channels : FL1 (GFP) and FL3 (RFP)). The accuracy of this measurement was first validated by comparison with manual countings with a hemocytometer. A volume of 40 μL of the mix (corresponding to 8 10^5^ cells) was then plated on 6 cm Petri dishes filled with 2 mL of 2% Phytagel (Sigma-Aldrich), following Dubravcic *et al* (Dubravcic et al., 2014). Cells were then incubated for 24 hours at 22°C. For each mix, three technical replicates were performed by plating three 40 μL droplets of cell suspension on 3 different Petri dishes.

### Quantification and statistical analysis

#### Quantification of spore bias in chrono-chimeras

24h after plating, spores were harvested by washing the three Petri dishes corresponding to the three technical replicates in 500 μL SorC buffer. Spores suspensions were incubated for 5 minutes with 0.5% Triton X-100 and then centrifuged for 7 min at 2000 rpm to remove stalk cells or unaggregated cells that would have survived the 24h-starvation-period. Finally, the pellet was resuspended in 800μL SorC buffer and the proportion of GFP and/or RFP-spores was scored using a cytometer. Spore bias of the focal population was quantified as the deviation between its proportion in the spores *f*_*S*_ and its proportion *f* in the initial mix.

To quantify the effect of growth phase heterogeneity on spore bias, we measured spore bias in chrono-chimeras composed of cells harvested in EE, ME, LE and ES phases of vegetative growth. Starting from the same batch of frozen aliquots, we tested a range (between 7 and 15 for each binary combination) of proportions *f* to assess frequency-dependent effects on spore bias. In principle, transformed strains were expected to produce no bias upon co-aggregation if the inserted fluorescent markers were strictly equivalent. However, we realized that chimeras composed of AX3-RFP and AX3-GFP cells grown in co-culture and both harvested in EE phase yielded a reproducible and significant bias (Fig. S2 A). The same bias was observed after having repeated the transformation protocol. In order to compensate for such labelling effect on spore bias, we substracted from our measures of the spore bias the bias predicted based on co-culture of differently labeled populations. Such intrinsic bias was computed for every frequency by interpolating with a third degree polynomial constrained with *f* = 0 and *f* = 1 (Fig. S2 A). In order to validate the use of this correction, we confirmed that spore biases are reversed when the fluorescent labels are swapped in a chrono-chimera where cells in EE and ME phase are mixed (Fig. S2 B).

#### Quantification of loner bias in chrono-chimeras

loner bias was estimated in chrono-chimeras composed of a comparable number of cells from the two populations (*i*.*e f ∼* 0.5). Chrono-chimeras were prepared as previously described and plated on a Petri dish that was scanned and imaged at regular time intervals (5 min). Images were taken with an automated inverted microscope Zeiss Axio Observer Z1 with a Camera Orca Flash 4.0 LT Hamamatsu, using a 10X objective, which yielded phase contrast and fluorescence images. Cell aggregation was considered complete when the last streams disappear. At that time, the number of unaggregated cells from the two populations was scored. Images corresponding to different areas of the Petri dish were first analysed using ImageJ software (Schindelin et al., 2012): aggregates were manually contoured and discarded and the “Find edges” ImageJ function was applied to highlight the contour of individual unaggregated cells. *f*_*L*_, the fraction of loners produced by the focal population, was then estimated on several images as the number of unaggregated cells from this population divided by the total number of unaggregated cells. Based on this observable we were able to quantify the bias in the fraction of unaggregated cells as the deviation between the proportion of cells from the focal population found in the pool of unaggregated cells (*f*_*L*_) and *f*.

The chrono-chimeras for which we measured the loner bias in parallel of the spore bias (Fig. 3) were started from a stock of frozen aliquots with higher initial cell density than that used to measure spore bias for a range of *f* values (Fig. 2). If spore bias variation is overall consistent, the EE/ES chimeras showed a more variable and mostly positive spore bias, suggesting possible long-term memory effects of population density.

#### Motility assay

Cells were first starved as previously described. After the last centrifugation, the pellet was re-suspended in 3 ml of buffer with a density of 10^4^ cells mL^*−*1^. This cell density was sufficiently low for cells not to touch with one another during the assay. The cell suspension was then poured in an empty 6 cm Petri dish. After 30 min –the time for the cells to attach to the bottom of the dish– cells trajectories were tracked for 1h (one image per 30 seconds) under an inverted microscope equipped with a moving stage and a 5X objective (alike to the measure of ‘loner bias’). A large area of the Petri dish was scanned to analyze around 600 cells trajectories per sample. Cell trajectories were then automatically extracted from the time lapse movies using the Python package *Trackpy* (Allan et al., 2018). Three biological replicates were imaged for populations harvested in EE, ME, LE and ES phase. Another script was used to analyse trajectories. Mean square displacement (MSD) was computed as a function of time lag for every single cell. The slope of the log(MSD) vs log(time lag) curve at low Δ*t* values (Δ*t <* 150 seconds) was used as a criterion to distinguish slowly moving (slope *<* threshold) from fast-moving cells (otherwise). The threshold value was set to 0.5 in order to separate the two modes of the slope distribution (Fig. S5), and the proportion of cells belonging to each class was scored. Individual cell total displacement was quantified as the sum of cells displacements between two successive frames along the trajectory.

#### Cell-substratum adhesion assay

Cell-substratum adhesion was quantified based on cells’ ability to attach to the bottom of a TC treated culture flask *(CytoOne, CC7682-4325)* as in Adiba et al. (2022). Cells were first starved as previously described. After the last centrifuging, the pellet was re-suspended in 10 ml buffer and cell density was adjusted to 2, 5.10^5^ cells mL^*−*1^. The cell suspension was then incubated in a 25 *cm*^2^ flask (*CytoOne* CC7682-4825) for 30 minutes at 22 °C, the time for cells to attach to the bottom of the flask. Each culture flask was gently shaken to resuspend cells that were not attached to the bottom of the flask. The supernatant (containing unattached cells) was transferred into a 15 *ml* tube. Cell density in the supernatant was measured using a hemocytometer. The fraction of adhesive cells was obtained by dividing the density of cells in the supernatant by the total cell density inoculated in the flask and used as a proxy for cell-substrate adhesion level. This assay was performed on three biological replicates for populations harvested in EE, ME, LE and ES phase.

#### Cell-cell adhesion assay

Cell-cell adhesion was quantified with a modification of the method by Gerrish (Gerisch, 1968). Cells were first starved as previously described. After the last centrifugation, the pellet was re-suspended in 0.5 ml buffer at a density of 10^6^ cells mL^*−*1^. The cell suspension was rotated at 150 rpm and 22°C for 1 hour, allowing cells to form multicellular clumps. The number of unaggregated cells (singlets and doublets) was determined using a hemocytometer. The percentage of cells that had been recruited into aggregates was calculated as the total cell density minus the density of unaggregated cells, divided by the total cell density. This quantity was used as a proxy for cell-cell adhesion level. The assay was performed on three biological replicates for populations harvested in EE, ME, LE and ES phase.

#### Statistical analysis

Significance of pairwise comparisons was established based on two-sample Student t test or Mann–Whitney U test using the python module *Scipy* (Virtanen et al., 2020). Significance of the effect of populations growth phase on cells’ total displacement was tested with a linear mixed effects model using replicates as random effects (‘nlme’ library, R). The significance level was set equal to 5%.

## Limitations of the study

A first limitation of our experimental approach comes from the high level of variability in spore bias measurements. Such variability seems to be irreducible even in controlled experimental conditions, and constrains the extent to which the effect of small phenotypic differences can be quantified. For this reason, we have pushed temporal differences in chronochimeras to extreme levels, which may not be attained in natural settings.

Second, our experimental setup does not allow to count the total number of loners produced at the end of aggregation, but only their composition. As a result, we could not test if the bias established during chronochimeras aggregation explains exactly the bias in the spores pool composition observed at the end of development (which would anyway require to assume a fixed proportioning of cell types in the multicellular body).

Finally, when exploring the mechanistic basis of loner bias, we did not explore single-cell heterogeneity in signalling and chemotaxis. Characterizing cAMP signaling dynamics of populations harvested in different growth phases would allow to test the hypothesis that variation in population signaling properties is a source of loner bias (Rossine et al., 2020). In particular, differences in signalling may explain the inconsistency between the loner bias observed in EE/ES chronochimeras and the motility properties of ES-cells.

## Author contributions

MF, SA and SD: conceptualization, methodology. MF: investigation. MF, SA and SD: writing—original draft and revised manuscript. SD: funding acquisition. All authors contributed to the article and approved the submitted version.

## Supplementary Information

**Figure S1:**
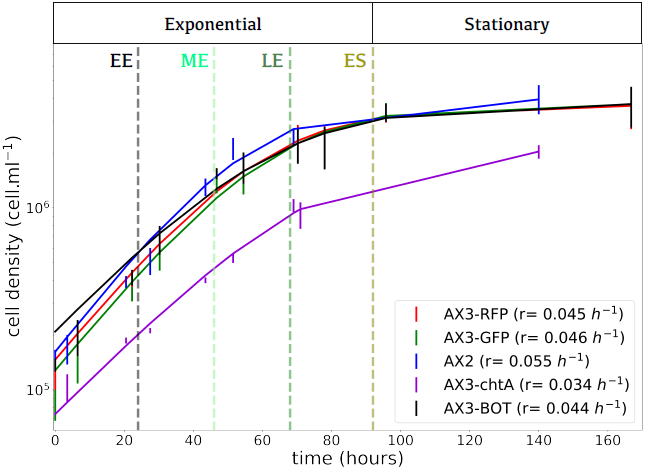
Growth kinetics of the different strains used in the experiments. Cell density of cultures grown in 25 *cm*^2^ flasks with 10 *ml* HL5 medium were assessed in triplicates using an hemocytometer. Before the beginning of the experiment, pre-cultures were prepared from frozen aliquots as described in the Methods. All strains had similar growth kinetics, but ChtA’s initial density was slightly lower due to fluctuations at the moment of dilution. The exponential and stationary phases of growth are indicated, as well as the times of harvesting considered in chronochimeras (vertical dotted lines), as explained in the methods. The average net growth rate in exponential phase (computed by linear interpolation between 20 and 50 hours) is reported for all the strains.

**Figure S2:**
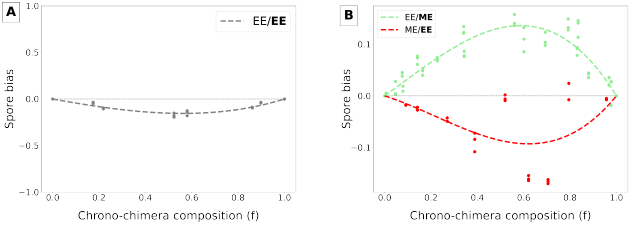
A: AX3-GFP spore bias at the end of chimeric development with AX3-RFP cells. The two populations were grown in co-cultures for 24 hours prior to aggregation to ensure they are in the same growth phase at the onset of the social cycle. Transformation with different plasmids introduced a spore bias (consistent when the transformation was repeated). We used this frequency-dependent bias to correct the measures realized in isogenic chronochimeras (main text and Methods). **B: Check for consistency of spore bias measures**. Corrected spore bias measured in two types of ‘chrono-chimeras’, using the AX3-GFP as the reference population: the AX3-GFP population was harvested in ME phase and mixed with a EE AX3-RFP population (light green line), and vice-versa (red line). Spore biases are reversed when the fluorescent labels are swapped, validating the use of the frequency correction.

**Figure S3:**
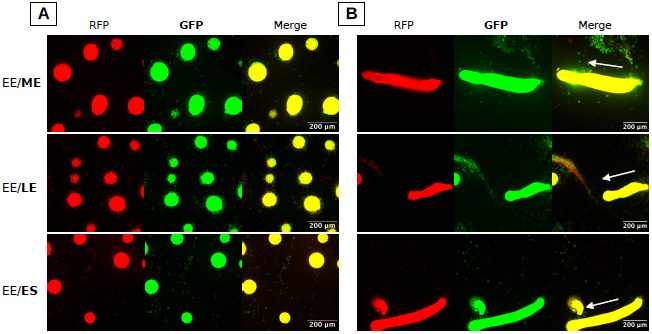
Heterogeneity in populations growth phase at the onset of starvation does not translate into detectable cell sorting during *Dictyostelium* social cycle. AX3-RFP cells harvested in EE phase of the growth cycle are mixed in chimeras with AX3-GFP cells harvested either in ME (first row), LE (second row) or ES phase (third row). **A:** RFP and GFP-cell populations do not segregate during aggregation but rather form chimeric aggregates with no noticeable difference in composition between aggregates, nor evident spatial sorting within aggregates. **B:** RFP and GFP-cell populations do not show significant signs of sorting along the slug axis during its migration. The white arrow indicates the direction of slug migration. Notice that since all cells bear a fluorescent marker, the high density of the mound and slug stages make it impossible to distinguish single cells in the images.

**Figure S4:**
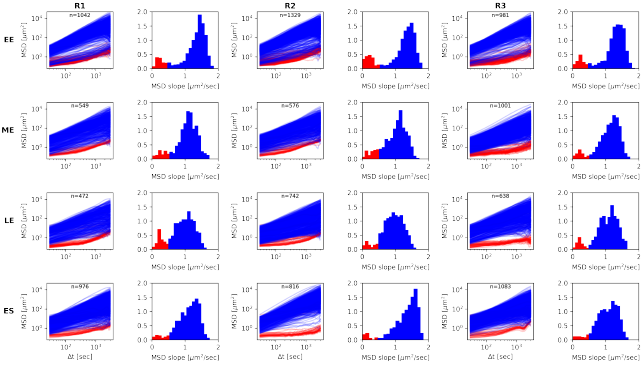
Bimodal distribution of single-cell motility depends on growth phase. Individual mean square displacement of cells from populations harvested in EE, ME, LE and ES phase as a function of time lag (Δ*t*). Cells were clustered into two classes based on the initial rate of increase of the MSD (slope of the log MSD vs log Δ*t*, Δ*t <* 150 secs, below or above the threshold value 0.5). As shown in Fig. 4 C, the proportion of cells belonging to the non-migrating class decreases in the course of vegetative growth.

**Figure S5:**
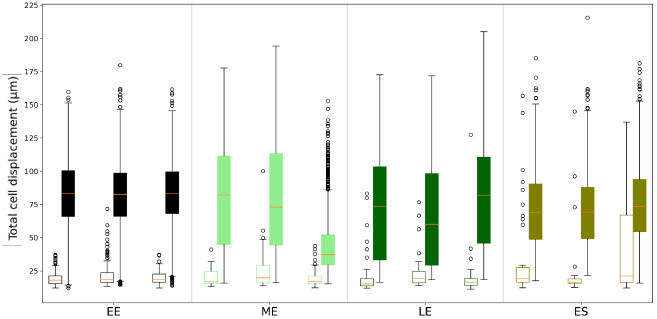
Total cell displacement of cells harvested from populations in EE, ME, LE and ES phases. Cells total displacement during the time of the experiment (1h) measured for the non-migrating cells class (empty boxes) and the migrating cells class (filled boxes) for populations harvested in EE, ME, LE and ES phase. Total displacement of non-migrating cells is significantly lower than that of migrating cells in every populations (Mann–Whitney U test, p-values *<* 0.0005). No significant variation in total displacement of migrating cells was observed between populations harvested in different growth phases (mixed effects model ANOVA, p-value=0.4691).

**Figure S6:**
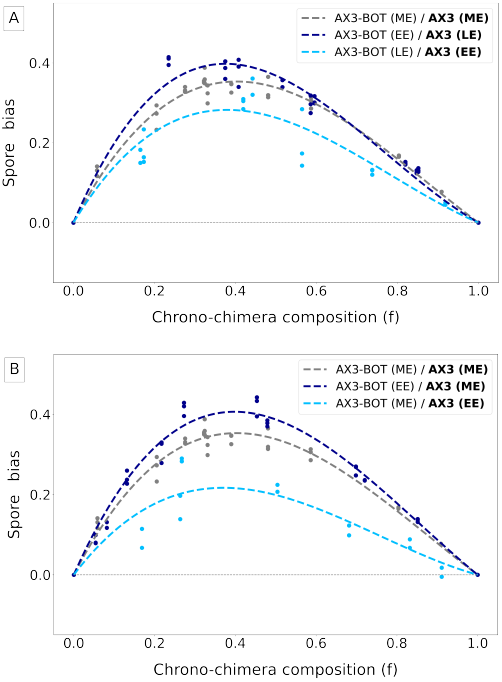
Spore bias measured in chrono-chimeras with a lineage selected for increased adhesion. Aggregation was initiated by mixing a strain that was obtained through experimental evolution for higher adhesiveness to the substratum (AX3-Bottom, Adiba et al. (2022)) with its ancestor (AX3). Three different combinations of growth phases are considered (as indicated in the legend). **A:** Strains are grown in co-culture and harvested in ME phase (gray line), the focal population is harvested in EE phase and the other in ME phase (dark blue line), or vice-versa (light blue line). **B:** Strains are grown in co-culture and harvested in ME phase (gray line), the focal population is harvested in EE phase and the other in LE phase (dark blue line), or vice-versa (light blue line). Spore bias variations are consistent whether the more advanced population is harvested in middle or late exponential phase, as it was observed for isogenic strains (Fig. 2), but starting from a high baseline bias.

**Figure S7:**
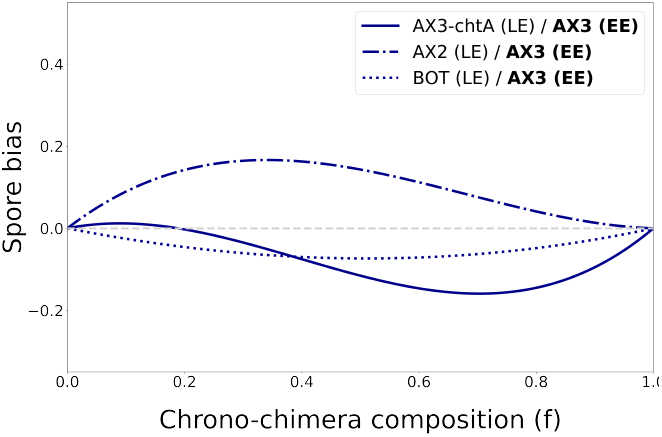
Contribution of growth phase differences to spore bias depends on genetic background. Comparison of spore bias when the focal population AX3 (EE) is mixed with different strains in late exponential phase (LE). For every chimera, the curve indicates the deviation of the fitted spore bias from that for reference chimeras, where the two strains were harvested in the same growth phase (ME). The qualitative effect of growth phase differences is not consistent across the different chrono-chimeras, suggesting that it does not simply add up to variation induced by genetic differences, but instead depends on the genetic background.

